# Structural changes in the gut microbiome of short-term and long-term medical workers compared to normal controls

**DOI:** 10.1101/772590

**Authors:** Ning Zheng, Shenghui Li, Bo Dong, Wen Sun, Huairui Li, Yongli Zhang, Peng Li, Zhiwei Fang, Changming Chen, Xiuyan Han, Bo Li, Siyi Zhang, Miao Xu, Guixin Zhang, Yi Xin, Yufang Ma, Xianyao Wan, Qiulong Yan

**Author notes:** Correspondence: Qiulong Yan & Xianyao Wan. These authors have contributed equally to this work (Ning Zheng, Shenghui Li, Bo Dong, Wen Sun).

## Abstract

**Objective:** Hospital environment has been implicated in enrichment and exchange of pathogens and antibiotic resistances, but its potential in shaping the symbiotic microbial community of the hospital staff is unclear. This study was designed to evaluate the alteration of gut microbiome in medical workers compared to non-medical controls.

**Design:** Prospective cross-sectional cohort study.

**Setting:** Intensive care unit (ICU) and other departments from a center in northeast China.

**Subjects:** 175 healthy medical workers (1-3 months short-term workers, n = 80; >1 year long-term workers, n = 95) and 80 healthy normal controls.

**Interventions:** None.

**Measurements and Main Results:** Fecal samples of all subjects were analyzed using the 16S rRNA gene sequencing. Medical workers exhibited remarkable deviation in gut microbial within-sample diversity and enterotypes stratification, and shift in overall microbial structure. Short-term workers were significantly more abundant in taxa including *Lactobacillus*, *Butyrivibrio*, *Clostridiaceae_Clostridium*, *Ruminococcus*, *Dialister*, *Bifidobacterium*, *Odoribacter* and *Desulfovibrio*, and with lower abundances of *Bacteroides* and *Blautia* compared with the controls. While long-term workers were enriched in taxa including *Dialister*, *Veillonella*, *Clostridiaceae_Clostridium*, *Bilophila*, *Desulfovibrio*, *Pseudomonas* and *Akkermansia*, with lower abundances of *Bacteroides* and *Coprococcus* compared with the controls. In addition, medical worker’s working years (short-term vs. long-term), hospital department (resident doctor vs. nursing staff) and work position (ICU vs. not-ICU) revealed considerable effects on their gut microbiome. Moreover, by analyzing the environmental samples (n = 9) around the inpatient wards and the hospital, we showed that the gut microbiota of medical workers was closer to environmental microbiota than that of the normal controls, probably in correlation to lasting exposure to the pathogenic taxa (e.g. *Pseudomonas*) in health workers.

**Conclusions:** Our findings demonstrated structural changes in the gut microbial community of the medical workers. Further studies are proposed for investigating the potentially physiological influence of the altered gut microbiome in medical participants.

**IMPORTANCE:** In this study, we for the first time focused on the influence of hospital environmental factors on gut microbiota of medical workers. The significance of our study is not limited to revealing the remodeling effect of the hospital environment on the gut microbiota of medical workers. Based on these, we also propose targeted and operational recommendations that can promote the health of hospital staff.

## INTRODUCTON

Unique chemical and physical conditions make the microbial community in the hospital environment vastly different from the external natural environment (1–3). Even in the same hospital, different regions usually have their own micro-ecological characteristics due to the differences in selection pressure. Poza *et al.* compared the microbial population in the entrance hall and intensive care unit (ICU) of a hospital in Spain, and found that the microbial community diversity in ICU is significantly lower than that in the former unit and the microbial composition in ICU is closer to the microbial populations of ICU-hospitalized patients (1). A prior study has analyzed the surface microbial flora of the medical ICU (MICU) and respiratory care center by using 16S rRNA gene sequencing, and found that the more stringent cleaning system in MICU can lead to a decrease in microbial community diversity and abundance of pathogens such as *Acinetobacter*, *Streptococcus* and *Pseudomonas* (4). Studies on gut microbial strains of preterm infants in the neonatal ICU revealed that almost all bacterial strains associated with infant gut colonization can be detected in the hospital room environment (5).

Microbial ecology in hospital has also been preliminary verified to impact on the health status of the hospital staffs (6). Scott Kelley *et al.* showed that the unique microbiological characteristics of the hospital environment promote the occurrence of pulmonary infections in staff (7). Subsequent studies revealed that the healthcare workers had higher incidence of *Clostridium difficile* infection (8) and may act as a potential vector of infectious diseases (9). The revolutionary advancements in microbial DNA sequencing technology have enabled us to study how microorganisms colonize and spread in specific environments. However, the composition of gut microbiota in medical workers and the influence from hospital environment are still unclear. Here, to investigate the characteristics of the gut microbiome in medical workers, we compared the microbial community composition of fecal samples obtained from 175 medical workers and 80 normal controls. We used 16S rRNA gene sequencing technology and bioinformatic analyses to identify microbial diversity, compositional and functional characteristics of the medical workers. Environmental samples (n = 9) of the hospital were also analyzed to evaluate its effect on the medical worker’s gut microbiota.

## MATERIALS AND METHODS

### Clinical trial and Ethics statement

This study received approval from the ethics committee of the First Affiliated Hospital of Dalian Medical University, and written informed consent was obtained from each participant. This study has been reported to Chinese Clinical Trial Registry/http://www.chictr.org (Identifiers: ChiCTR19000251156). The methods were carried out in accordance with the approved guidelines.

### Participants and sample collection

The First Affiliated Hospital of Dalian Medical University own more than 4,000 employed doctors and staffs, approximately 3,700 beds and nearly 100,000 inpatients, with an annual service volume of more than 2.5 million people. In this study, 175 full-time medical workers, comprising 80 short-term workers who had entered the hospital for 1-3 months and 95 long-term workers for at least 1 year, were recruited. 49.7% of the medical workers were from the ICU, which had 20 beds and treats more than 1,000 critically ill patients each year. The non-medical control group included 80 healthy individuals who visited the same hospital for physical examination. Subjects of short-term workers and normal controls had not worked at other hospitals in last half year before participating in this study.

Subjects were excluded if they had symptoms of respiratory or digestive tract infections, or if they were treated with antibiotics or anti-inflammatory agents in recent two months prior to sampling. Subjects with severe obesity [body mass index (BMI) ≥32 kg/m^2^], diabetes, metabolic syndrome, inflammatory bowel disease or severe cardiovascular diseases (such as coronary artery disease or stroke) were also excluded. All participants in this study were urban residents who lived in the city of Dalian, China.

#### Fecal sample collection

Fresh fecal samples were collected at the hospital from each subject and stored at a −80°C freezer within 0.5 hours. Individuals with diarrhea, constipation, abnormal fecal color, viscous stool (fat content over high) were excluded.

#### Environmental sample collection

Several environmental samples, including ventilation system dust (air, n = 3), sewage (n = 3) and soil (n = 3), were collected around the inpatient wards and the hospital. The dust of exhaust port was collected by sterile cotton swab at the ICU (n = 2) and other wards (n = 1). Soil samples were collected approximately 10 cm^2^ (per sampling site) within 20 meters of the building which contained the ICU ward. One-liter bilge water from sewage outlet was collected at sewage treatment tank of this building. Each environmental sample was pooled from 3-5 sampling sites. Transport samples in an insulated transport container with frozen gel packs and dry ice to keep the samples cold. All samples stored at −80°C prior to DNA extraction and chemical analysis.

### DNA preparation and sequencing

Total bacterial genomic DNA were extracted from about 220 mg of feces or environmental samples (the pellets of water samples were collected after centrifugation) using the QIAamp DNA Stool Mini Kit (Qiagen, Germany) following the manufacturer’s instruction. The final DNA concentration and purification were determined by NanoDrop 2000 UV-vis spectrophotometer (Thermo Scientific, Wilmington, USA). The V3-V4 hypervariable regions of the bacterial 16S rRNA gene were amplified with primers 338F (5’-ACTCCTACGGGAGGCAGCAG-3’) and 806R (5’-GGACTACHVGGGTWTCTAAT-3’) by thermocycler PCR system (GeneAmp 9700, ABI, USA). The PCR reactions were conducted using the following program: 3 min of denaturation at 95 °C, 27 cycles of 30 s at 95 °C, 30s for annealing at 55 °C, and 45s for elongation at 72 °C, and a final extension at 72 °C for 10 min. PCR reactions were performed in triplicate 20 μL mixture containing 4 μL of 5 × FastPfu Buffer, 2 μL of 2.5 mM dNTPs, 0.8 μL of each primer (5 μM), 0.4 μL of FastPfu Polymerase and 10 ng of template DNA. The resulted PCR products were extracted from a 2% agarose gel and further purified using the AxyPrep DNA Gel Extraction Kit (Axygen Biosciences, Union City, CA, USA) and quantified using QuantiFluor™ -ST (Promega, USA) according to the manufacturer’s protocol.

Purified amplicons were pooled in equimolar and paired-end sequenced (250bp PE) on an Illumina HiSeq2500 platform according to the standard protocols by Majorbio BioTech Co. Ltd. (Shanghai, China).

### Bioinformatic analyses

Raw sequencing reads were eliminated from analysis if they contain >8 homopolymers, >2 mismatches in the primers, or >1 mismatches in the barcode sequences. High-quality paired-end sequencing reads were then analyzed under the quantitative insights into microbial ecology (QIIME2, https://qiime2.org/) platform (10) and the standard tools/plugins provided by QIIME2. Briefly, 16S sequences were performed for quality control and to feature table construction using the DADA2 algorithm (11). Possible phiX reads and chimeric sequences were removed, and the remaining reads were truncated from 0 to 260 bases (for both forward and reverse reads) to avoid the sequencing errors at the end of the reads. Paired-end reads were overlapped at the maximum mismatch parameter of 6 bases, which means a minimum similarity of approximately 90% on the overlap region of the forward and reverse reads. The representative sequences (named “feature” in QIIME2 nomenclature) were then generated by removing the redundant and low occurrence (n < 5 in pool samples) sequences. We used the term “operational taxonomic unit (OTU)” instead of “feature” in the whole article for convenience. Phylogenetic analysis of OTUs were realized via the q2-phylogeny plugin, and taxonomic assignment of the OTUs were determined via a pre-trained Naive Bayes classifier (trained on the Greengenes 13_8 99% OTUs (12)) using the q2-feature-classifier plugin. To avoid the depth deviation of samples potentially biased in community diversity and other analyses (13), 22,000 reads were randomly selected from each sample when calculating the OTU relative abundance.

Four estimators of the alpha diversity of microbial community, including Shannon index, observed OTUs, Faith’s phylogenetic diversity and Pielou’s evenness, were used in this study and calculated using the QIIME2 q2-diversity plugin. Bray-Curtis dissimilarity was used to evaluate between-sample (beta) diversity, which was calculated based on the *vegan* package in R platform.

Enterotype of the fecal samples was determined based on genus level microbial community composition using a reference-based alignment algorithm (http://enterotypes.org/) (14).

Functional composition of the samples was generated using the PICRUSt2 algorithm (15). For each sample, the composition of Kyoto encyclopedia of genes and genomes (KEGG (16)) orthologs (KOs) was predicted based on the functional information of the reference OTUs. KEGG modules and pathways composition was generated according to the assignment of KOs at https://www.kegg.jp/. For each pathway in differential analysis, the reporter score was arithmetic averaged from the Z-scores of individual KOs (calculated via Mann–Whitney U test and adjusted for multiple testing using the Benjamin-Hochberg procedure) (17, 18).

The raw sequencing dataset acquired in this study have been deposited to the European Bioinformatics Institute (EBI) database under the accession code PRJEB28135.

### Statistical analyses

Linear discriminant analysis effect size (LEfSe) analysis (19) was performed on normalized taxa composition using web server (http://huttenhower.sph.harvard.edu/galaxy). Distance-based redundancy analysis (dbRDA) was performed on taxa composition and functional profiles with R *vegan* package, according to the Bray-Curtis dissimilarity, and visualized with R *ade4* package. To access the effect of host properties on the gut microbiome, permutational multivariate analysis of variance (PERMANOVA) was implemented with *adonis* function of the *vegan* package. *P*-value < 0.05 was considered statistical significant, and the *q*-value was calculated to evaluate the false discovery rate (FDR) for correction of multiple comparisons.

## RESULTS

### Participant characteristics and sequencing coverage

This study involved 255 healthy Chinese participants, including 175 medical workers and 80 normal controls (NC group). Basic characteristics of all participants are summarized in **Table 1**, and the detailed information of all participants are shown in **Table S1** (Supplemental Digital Content 1). The medical worker population consisted of 80 short-term workers (ST group) who had worked in the hospital for 1-3 months and 95 long-term workers (LT group) working in the hospital for at least one year. Medical workers and normal controls were matched in gender, age and BMI. 66% of the medical workers and 61% of the normal controls were female. 54% of the medical workers were resident doctors, and the remaining were nursing staff. 50% of the medical workers were working in the ICU.

**Table 1.** Characteristics of participants.

| Characteristics | Medical workers |  | Normal controls | *P*-value (ST vs. NC) | *P*-value (LT vs. NC) |
| --- | --- | --- | --- | --- | --- |
|  | Short-term | Long-term |  |  |  |
|  | (ST) | (LT) |  |  |  |
| No. of individuals | 80 | 95 | 80 |  |  |
| Gender (F/M) | 52/28 | 64/31 | 49/31 | 0.743 | 0.43 |
| Age, years | 30.7±6.7 | 31.3±6.4 | 32.7±8.6 | 0.093 | 0.203 |
| BMI, kg/m<sup>2</sup> | 22.1±3.5 | 22.7±3 | 22.6±2.6 | 0.309 | 0.81 |
| Working period | 1-3 months | >1 year |  |  |  |
| Position (doctor/nurse) | 40/40 | 54/41 |  |  |  |
| Department (ICU/not) | 38/42 | 49/46 |  |  |  |

*P*-value was calculated using Fisher's exact test (for gender) and Student's *t*-test (for age and BMI). BMI, body mass index. ICU, intensive care unit.

We identified a total of 54,812 OTUs for bacterial level phylotypes: 73.9% of these OTUs were robustly classified into genus-level taxa, accounting for 80.3% of the total reads; 91.1% of the OTUs could be classified into the level of specific families, accounting for 95.5% of the total reads.

### Distinct gut microbial diversity and structure in medical workers

We performed the PERMANOVA analysis on the gut microbial composition of the entire cohort, to test if the status of medical work was associated with the holistic community structure. Stratification of medical workers and normal controls accounted for 4.6% (adonis *P* < 0.001) of the gut microbiota variance. This effect size was relatively larger than other collected confounding factors, including the participants’ intrinsic parameters (gender, age and BMI), lifestyle and diet patterns (**Table S2**, Supplemental Digital Content 1), indicating that the status of medical work was one of the main reasons in shaping the gut microbiome in our cohort.

To investigate whether the gut microbial diversity and structure are differed between medical workers and normal controls, first, we assessed the microbial (alpha) diversity within each individual. The ST group showed the highest level of microbial diversity compared with the LT and NC groups, which was reflected in Shannon index, phylogenetic diversity and Pielou’s evenness (**Figure 1A**); although the LT group had significant increases in phylogenetic diversity and observed OTUs as contrasted to control subjects. Next, we performed enterotypes classification of all individuals and found that the ST group exhibited a significant deviation of enterotypes compared with the NC group (**Figure 1B**). Lastly, multivariate analysis (dbRDA) based on Bray-Curtis distance between microbial genera revealed remarkable differences among three groups (**Figure 1C**). Both the ST and LT groups were deviated from the NC group on the primary constrained axis (CAP1, explaining 5.3% of total variance) of the dbRDA plot, while the ST group was more distant. In addition, comparison of pairwise Bray-Curtis dissimilarity also revealed a similar fashion between medical workers and normal controls (**Figure 1D**), yet the gut microbiota of ST workers resembled more close to the controls’ gut microbiota than that of LT workers. Taken together, these findings revealed remarkable differences in gut microbial diversity and structure in medical workers.

**Figure 1.**
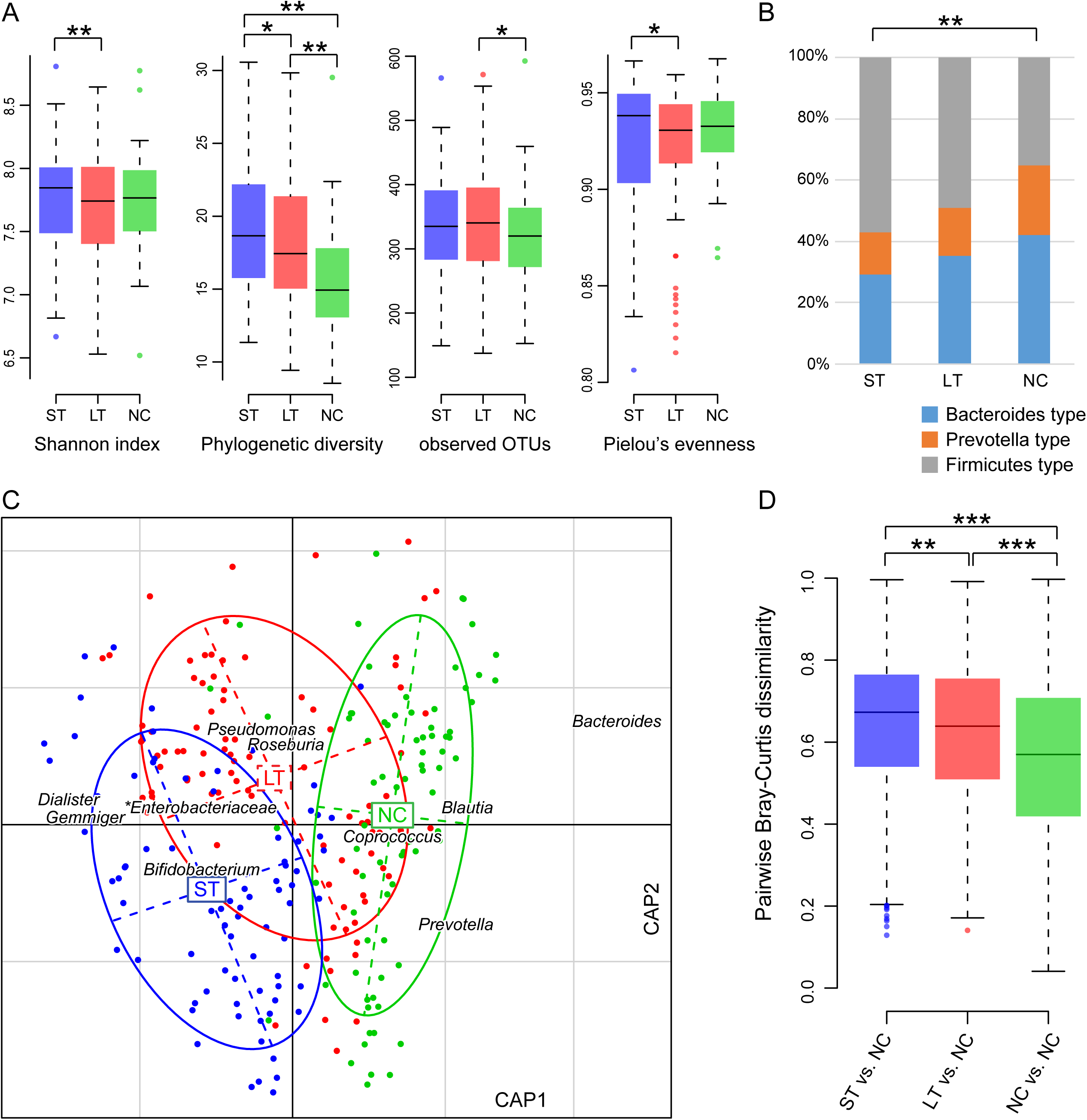
Alteration of gut microbiome diversity and structure in medical workers compared with normal controls. **(A),** Difference of within-sample diversity between medical workers and controls. Boxes represent the interquartile range (IQR) between first and third quartiles and the line inside represents the median. The whiskers denote the lowest and highest values within 1.5 times of IQR from the first and third quartiles, respectively. The dots represent outliers beyond the whiskers. *, *P* < 0.05; **, *P* < 0.01; Students’ *t* test. **(B),** Enterotypes stratification of the samples. **, *P* < 0.01; Kruskal-Wallis rank sum test. **(C),** dbRDA based on the Bray-Curtis dissimilarity between microbial genera, revealing significant deviation between medical workers and normal controls. The primary and second constrained axes (CAP1 and CAP2) of dbRDA plot are shown. Genera (square) as the main contributors are plotted by their loadings in these two axes. Lines connect samples in the same group, and circles cover samples near the center of gravity for each group. **(D),** Boxplot shows the pairwise Bray-Curtis dissimilarity between medical workers and normal controls. **, *P* < 0.01; ***, *P* < 0.001; Students’ *t* test.

### Phylogenetic alteration of gut microbiota

At the phylum level, Firmicutes, Bacteroidetes, Proteobacteria and Actinobacteria comprised the vast majority of the gut microbiota of all participants. Of these, Firmicutes was more abundant in the ST (average relative abundance, 66.5%) and LT groups (59.7%) than the NC group (55.8%), while Bacteroidetes was less abundant (average relative abundance in ST, LT and NC groups, 22.1%, 27.2% and 33.6%, respectively). Proteobacteria was slightly but not significantly more abundant in medical workers (6.1%/7.5% in ST/LT vs. 4.9% in NC).

To further explore signatures of the gut microbiome in medical workers and normal controls, we performed the LEfSe analysis on the microbial composition at all taxonomic levels. We identified 79 microbial taxa (57 ST-enriched and 22 NC-enriched) that significantly differed in relative abundance between the ST and NC groups (**Figure 2A**; **Table S3**, Supplemental Digital Content 1), and 59 microbial taxa (46 LT-enriched and 13 NC-enriched) that significantly differed between the LT and NC groups (**Figure 2B**). Compared with the normal controls, the ST group showed higher abundance in taxa belonging to Firmicutes (including *Lactobacillus*, *Clostridiaceae_Clostridium*, *Butyrivibrio*, *Ruminococcus* and *Dialister*), Actinobacteria (including *Bifidobacterium* and *Actinomyces*), *Odoribacter* and *Desulfovibrio*, and lower abundances of *Bacteroides*, *Blautia*, *Erysipelotrichaceae_Clostridium* and *Pseudomonas*. Compared with the normal controls, the LT group exhibited higher abundance of Firmicutes members (including *Clostridiaceae_Clostridium*, *Dialister* and *Veillonella*), *Bilophila*, *Desulfovibrio*, *Pseudomonas* and *Akkermansia*, and lower abundances of *Bacteroides* and *Coprococcus*.

**Figure 2.**
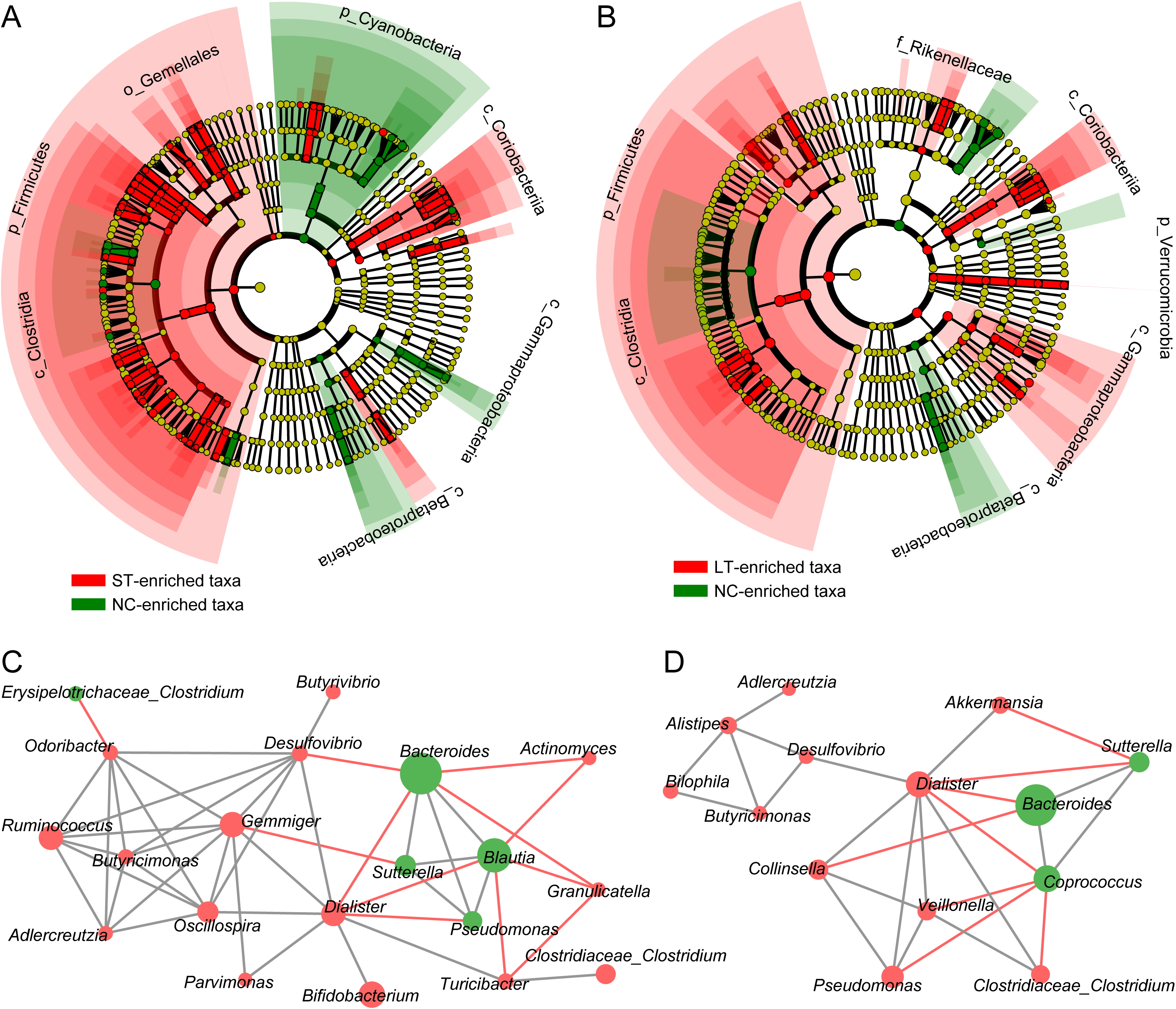
Medical worker-associated microbial taxa identified via LEfSe analysis. **(A-B),** Cladogram plots show the microbial taxa that significantly differed in relative abundance (LDA score > 2) between LT **(A)**, ST workers **(B)** and normal controls. Detailed information of the medical worker-associated taxa was shown at Table S2 (Supplemental Digital Content 1). **(C-D),** Co-occurrence networks show the interconnection of the ST worker-associated **(C)** and LT worker-associated **(D)** genera. Nodes depict medical worker-enriched (represented by red) and control-enriched (represented by green) genera with their taxonomic assignment displayed in the center. The size of the nodes indicates their average relative abundance in samples. Connecting lines represent Spearman correlation coefficient rho > 0.40 (represented by gray line) or < −0.40 (represented by red line).

Moreover, we performed co-occurrence network analysis and found a large number of interconnections within the ST-altered and LT-altered taxa, suggesting that these medical worker-associated taxa did not occur independently and interacted with the other taxa in its environment. On the ST-altered network, *Gemmiger*, *Butyricimonas*, *Desulfovibrio*, *Dialister*, *Bacteroides* and *Blautia* were keystone genera (**Figure 2C**), while on the LT-altered network, *Dialister*, *Veillonella* and *Coprococcus* were keystone genera (**Figure 2D**); these taxa might play essential roles in shaping the gut microbiota of medical workers.

### Functional characterization of gut microbiome

To characterize the functional capacity of the gut microbiome in medical workers, first, we observed a significant alteration of KEGG orthologue (KO) profiles in both ST and LT groups compared with normal controls (*adonis P* < 0.001; **Figure S1A**, Supplemental Digital Content 2). When compared with normal controls on the KEGG pathway, the gut microbiome of ST group was significantly more abundant in cell motility (mainly consisting of bacterial motility proteins and flagellar assembly), prokaryotic cellular community (biofilm formation), membrane transport (phosphotransferase system, secretion system and transporters), and xenobiotics biodegradation and metabolism (referring to degradation of multiple aromatic compounds and drugs) (all with reporter Z-score > 1.65; **Figure S1B-C**, Supplemental Digital Content 2). Similarly, the gut microbiome of LT group also revealed consistent tendency of these pathways, despite that only the functional catalogue of xenobiotics biodegradation and metabolism was significantly enriched.

### Effect of hospital department and work position

To assess the possible impact of host properties on gut microbiota, we tested if the hospital department (ICU vs. not-ICU) and work position (resident doctor vs. nursing staff) of the medical workers were associated with their gut microbiota. For both the ST and LT groups, significant deviation of gut microbial composition was observed in individuals with different hospital department and work position (**Figure 3A-B**). Compared with the not-ICU workers, workers in the ICU were with a significant increase of *Dialister*, Enterobacteriaceae, *Phascolarctobacterium*, *Pseudomonas*, *Veillonella* and *Streptococcus*, and a remarkable depletion of *Faecalibacterium*, *Blautia* and *Coprococcus* (**Figure 3C**). Compared with the nursing staffs, the gut microbiome of the resident doctors was with a significant increase of *Erysipelotrichaceae_Clostridium* and decrease of *Bacteroides*, *Blautia* and *Ruminococcus* (**Figure 3D**).

**Figure 3.**
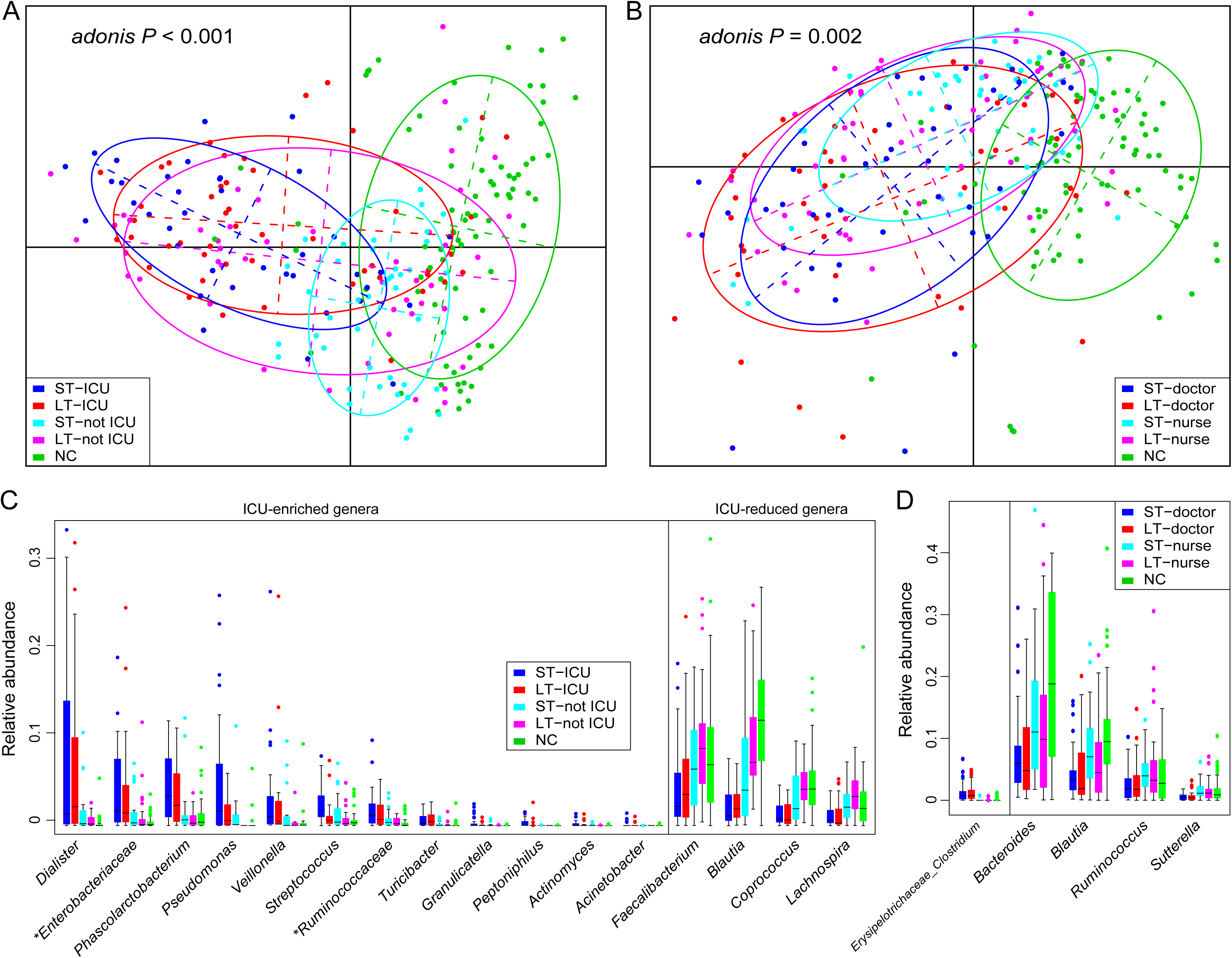
Differences in microbial composition of medical workers with different host properties. **(A-B),** dbRDA plots based on Bray–Curtis dissimilarity between microbial genera show observable deviation between different hospital departments **(A)** and work positions **(B)**. Points depict the samples, and color lines connect samples in the same group. **(C-D),** Boxplots show the significantly differed genera among different hospital departments **(C)** and work positions **(D)** in medical workers. Genera with *q* < 0.05 (Mann-Whitney U-test corrected by FDR) are shown.

### Effect of hospital environment on the gut microbiota of medical workers

To assess whether changes in the gut microbiota of medical workers were associated with their hospital environmental contact, we examined the microbial community composition of the hospital ecosystem via three representative environments: ventilation system dust (n = 3), sewage (n = 3) and soil (n = 3). Compared to the human gut microbiota, the environmental samples exhibited distinct microbial phylogenetic profiles (**Figure 4A**). Gut microbiota of both the ST and LT workers showed significantly lower Bray-Curtis dissimilarity to the environment samples compared to the normal controls (**Figure 4B**). Importantly, a genus which dominated in the environment samples, *Pseudomonas*, was significantly more abundant in the LT group than the ST and NC groups (**Figure 4C**). These findings suggested that working in the hospital was correlated with alterations in the gut microbiota of the medical workers.

**Figure 4.**
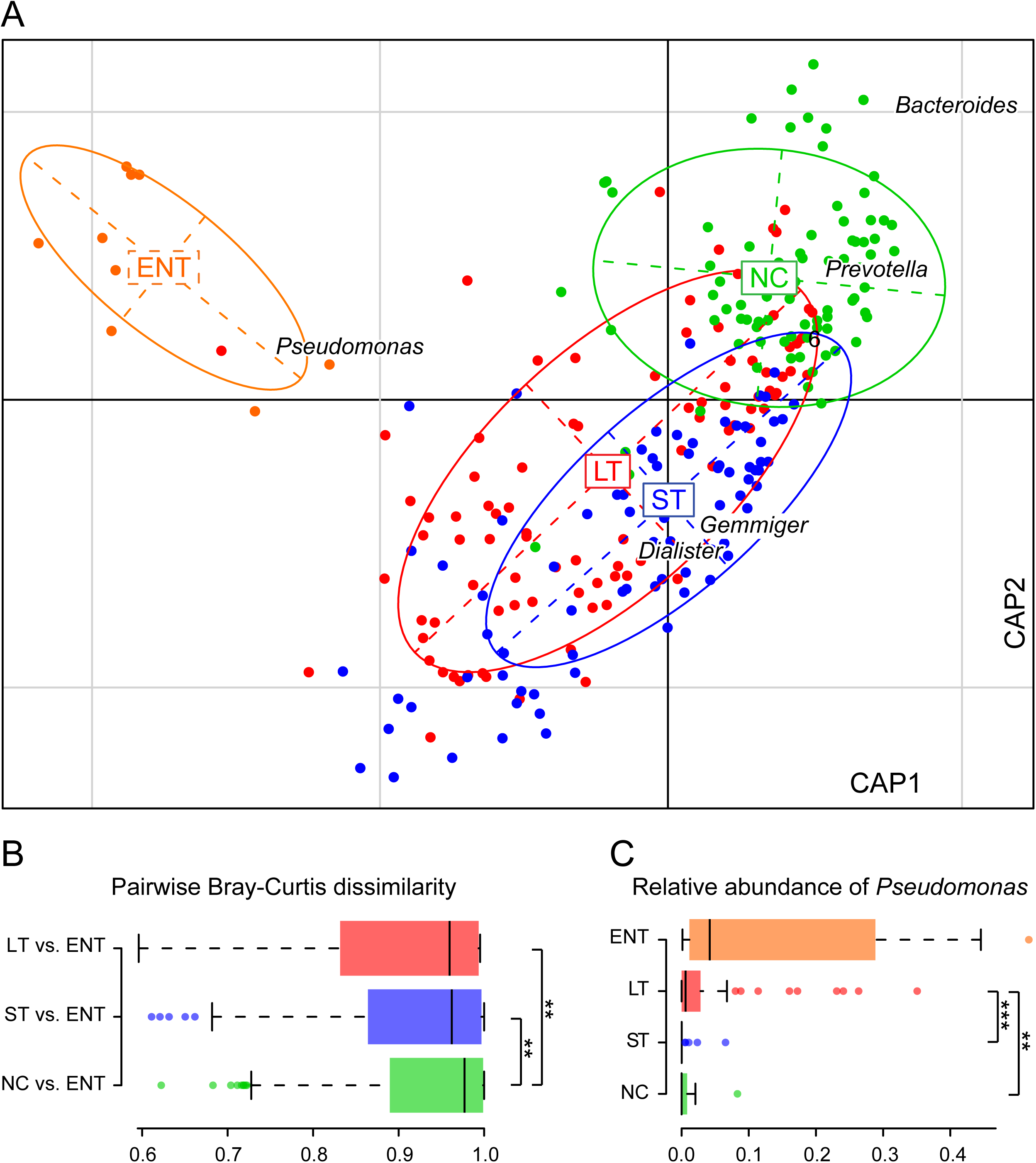
Effect of hospital environment on the gut microbiota of medical workers. **(A),** dbRDA plots based on Bray–Curtis dissimilarity between microbial genera show observable deviation between different environmental and human fecal samples. Points depict the samples, and color lines connect samples in the same group. **(B),** Boxplot shows the pairwise Bray-Curtis dissimilarity between human fecal samples and hospital environment. **(C),** Boxplot shows the relative abundance of *Pseudomonas* in environmental and human fecal samples. **, *P* < 0.01; ***, *P* < 0.001; Students’ *t* test.

## DISCUSSION

Medical workers are continually in contact with patients and exposed to the hospital environment in their daily work, however, the effect of such high-risk exposure on the gut microbiome of medical workers are rarely investigated. In this study, using 16S rRNA gene sequencing of the fecal samples, we demonstrated that the community diverisity, structure and function of gut microbiota of both short-term and long-term medical workers remarkably differed from that of non-medical controls. Medical worker’s working years (short-term vs. long-term), hospital department and work position showed nonnegligible effect on their gut microbiome. Moreover, alterations in the gut microbiome of medical workers were also in correlation to the environmental microbiome around the hospital.

Studying of hand hygiene provided effective approaches for infection prevention of medical workers (20). As an important evidence, previous studies had shown that the hand microbiome of healthcare worker can mediate the carriage of various infectious pathogens (21, 22). Gut microbiota, on the other side, also played an essential role in control of bacterial infection (23, 24). Thus, the altered gut microbiome in medical workers may be another risk factor in correlation to high level hospital-associated infections of these workers.

Compared with the normal controls, the ST workers’ gut microbiome showed significant enrichment of Firmicutes and reduction of Bacteroidetes. Firmicutes and Bacteroidetes maintain the major nutrient metabolic function of human gut (25), and the balance of Firmicutes/Bacteroidetes is associated with obesity (26), age (27) and metabolic dysfunctions (28). Several Firmicutes members, including *Lactobacillus*, *Butyrivibrio*, *Ruminococcus*, *Clostridiaceae_Clostridium* and *Dialister*, as well as the genera *Bifidobacterium*, *Odoribacter* and *Desulfovibrio* were enriched in the ST gut microbiome. *Lactobacillus* and *Bifidobacterium* strains are the main edible probiotics (29–31), while *Ruminococcus* is an important component of the human gut microbiome and yet drives the one of three enterotypes (32). Functionally, the gut microbiome of ST medical workers exhibited higher level of cell motility, prokaryotic cellular community, membrane transport, and xenobiotics biodegradation and metabolism compared to normal controls. Enrichment of these microbial functions is often related to metabolic disorders such as type 2 diabetes (33) and hypertension (34). Based on these findings, however, whether the ST workers’ gut microbiome that exert beneficial or pathogenic effects on the health of the host is still unclear.

Gut microbiome of the LT workers were enriched in genera including *Dialister*, *Veillonella*, *Bilophila*, *Alistipes*, *Clostridiaceae_Clostridium*, *Desulfovibrio*, *Pseudomonas* and *Akkermansia* compared with that of the normal controls. Enrichment of *Veillonella* in the gut microbiota are associated with liver cirrhosis (35) and cardiovascular disease (36), while the *Bilophila* strains can promote higher inflammation and induce metabolic dysfunctions via producing hydrogen sulfide (37, 38). Noticeably, the LT group showed a depletion of *Coprococcus*, a potential “anti-inflammatory” butyrate-producing bacteria in healthy human gut (39). Depletion of gut *Coprococcus* was correlated with lower quality of life (40) and mental diseases such as depression and chronic fatigue syndrome (28, 41). Collectively, these findings suggested potential unhealthy variation in the gut microbiota of LT workers; however, further investigations are in need to test this hypothesis.

The gut microbiome of ST group was more deviated from the normal controls than that of LT group, as revealed by both within-sample diverisity and bacterial compositional analysis (**Figure 1**). We speculated that the gut microbiota of medical workers underwent considerable changes at the initial stage of entering the hospital environment and could recover to a certain extent after their gradual adaptation to the working environment. In agreement with previous reports showing the remarkable and recoverable effect of dietary (42), lifestyle (43) and antibiotic usage (44) on human gut microbiota, our finding suggested the remodeling effect of environment on gut microbiota and the long-term resilience of gut microbiota after environmental change. In this study, alteration of gut microbiome in hospital can be attributed to comprehensive factors, of which we considered that the psychological factor is one possible: highly stressful work at the hospital for a medical newcomer often leads to anxiety, and such mental factors can affect gut microbiota (45, 46).

Affected by physical and chemical factors, different regions in the same hospital often have different ecological characteristics. In this study, nearly half of subjects were from the ICU, a medical unit for treating critically ill patients that have most rigorous equipment disinfection and daily cleaning. However, we found that, the gut microbiome of ICU workers (for both ST and LT groups) was more deviated from the controls than that of non-ICU workers, represented by a significant increase of *Dialister*, Enterobacteriaceae, *Phascolarctobacterium*, *Pseudomonas*, *Veillonella* and *Streptococcus*. Members of Enterobacteriaceae are among the most common pathogens in ICU (47), and their enrichment in medical workers might be attributable to the ICU environment. Notably, critically ill patient is the major source of environmental microbes in ICU and their gut microbiota is characterized by low abundance of key symbiotic bacteria (e.g. *Faecalibacterium prausnitzii* and *Ruminococcus*) and overgrowth of pathogenic bacteria (e.g. *Pseudomonas*, *Staphylococcus* and Enterobacteriaceae) (48, 49). Medical workers inevitably have contact with patients during routine treatment and nursing care, thereby increasing the risk of transmission of pathogenic bacteria (50).

In regards to the hospital environment, we showed that the environmental samples, including the air, sewage and soil, exhibited distinct microbial phylogenetic profiles compared to the human gut microbiome. As expected, gut microbiome of the both ST and LT medical workers showed significantly closer relationship to the environmental samples compared to that of the normal controls. The environmental samples were dominated by *Pseudomonas*, a ubiquitous pathogen that causes opportunistic nosocomial infection (51). This phenomenon is in agreement with the enrichment of *Pseudomonas* in the LT gut microbiome, supporting a potential transmission of clinical pathogen from the hospital environment to the long-term workers. Clearly, considering that current 16S rRNA sequencing were not fully capable of determining the transmission relationship of microbes among different habitats, further investigations based on new technologies (e.g. whole-metagenome shotgun sequencing) will lead to a better understanding of the mutual interaction.

This study has several limitations. 1) The study design was not longitudinal; we could not provide evidence for investigating the temporal alteration of the medical worker’s gut microbiome. 2) Despite the observation of differed gut microbiome in the medical workers compared to the normal controls, we could neither decipher the causality relationship of such differences, nor evaluate the influence of such difference on individual’s health. 3) The sampling sites and time points of the hospital environment is rare. Moreover, we didn’t collect the skin or respiratory tract samples of the medical workers which were more likely to be affected by the environment.

In this study, we for the first time focused on the distinct features of gut microbiota of healthy medical workers. The significance of our study is not limited to suggesting the remodeling effect of the hospital environment on the gut microbiota of medical workers. Based on these, we also propose targeted and operational recommendations that may promote the health of hospital staff, such as 1) relieving the anxiety of young medical workers who first enter the hospital, 2) regularly monitoring the gut microbiota of hospital staff, especially ICU-related workers, and 3) improving their gut micro-ecology by supplementing probiotics or prebiotics.

## Author contributions

Conceptualization and project administration, Ning Zheng, Shenghui Li, Xianyao Wan and Qiulong Yan;

Resources, Wen Sun, Yongli Zhang, Guixin Zhang and Xianyao Wan; Investigation, Ning Zheng, Bo Dong, Huairui Li, Changming Chen, Xiuyan Han, Bo Li and Siyi Zhang; Data curation and analysis, Shenghui Li, Bo Dong, Wen Sun, Zhiwei Fang, Peng Li, Changming Chen, Miao Xu and Guixin Zhang; Writing – original draft, Shenghui Li, Ning Zheng and Qiulong Yan; Writing – review & editing, Peng Li, Bo Dong, Wen Sun, Yi Xin, Yufang Ma and Xianyao Wan; Funding acquisition, Qiulong Yan and Xianyao Wan.

---

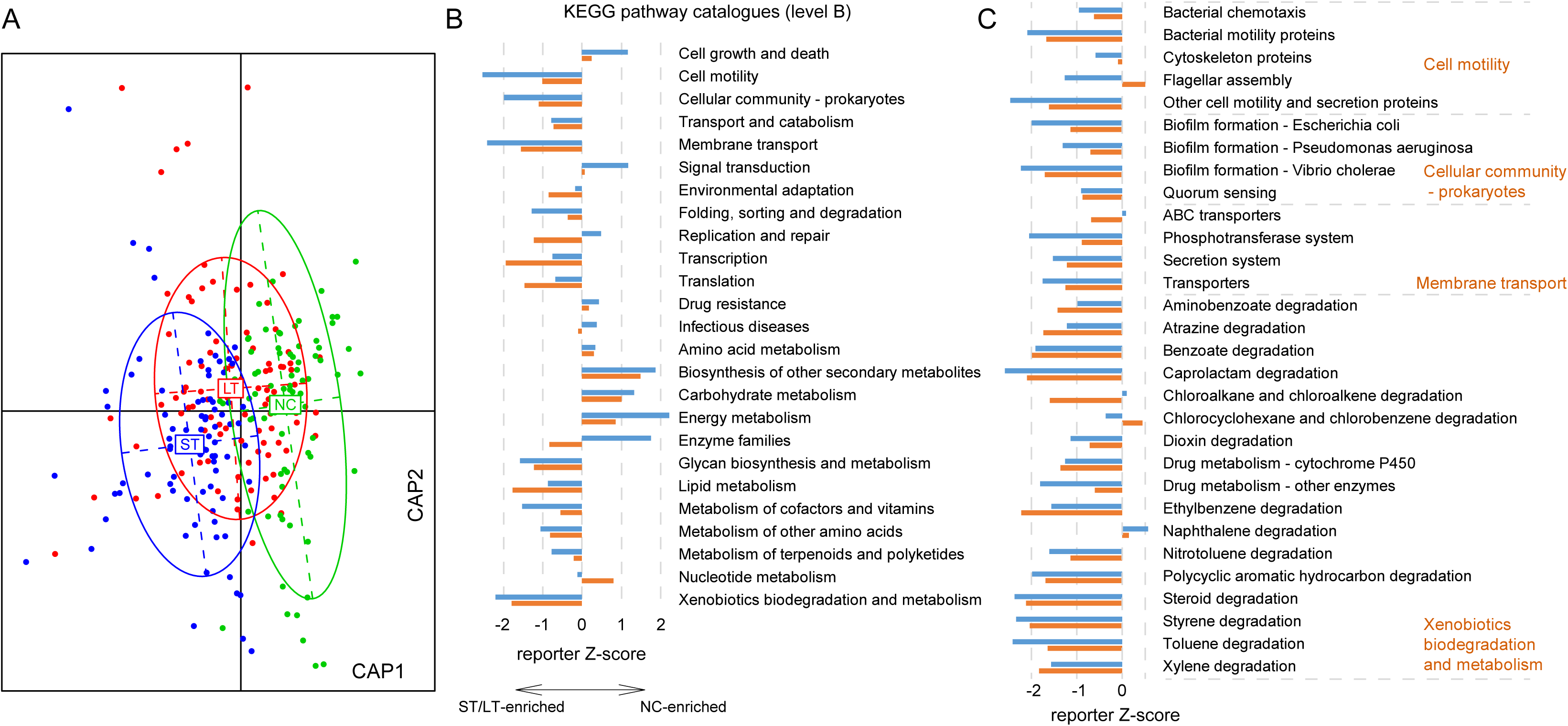

